# Both DNA Polymerases δ and ε Contact Active and Stalled Replication Forks differently

**DOI:** 10.1101/170795

**Authors:** Chuanhe Yu, Haiyun Gan, Zhiguo Zhang

**Affiliations:** Department of Biochemistry and Molecular Biology, Mayo Clinic, Rochester, MN, 55905, USA; Institute for Cancer Genetics, Department of Pediatrics, Department of Genetics and Development, Columbia University, 1130 St. Nicholas Avenue, Irving Cancer Research Center ICRC 407A, New York, NY10032

**Keywords:** DNA replication, replication stress, DNA polymerase, ChIP-ssSeq, strand specific sequencing

## Abstract

Three DNA polymerases (Pol α, Pol δ, and Pol ε) are responsible for eukaryotic genome duplication. When DNA replication stress is encountered, DNA synthesis stalls until the stress is ameliorated. However, it is not known whether there is a difference in the association of each polymerase with active and stalled replication forks. Here, we show that each DNA polymerase has distinct patterns of association with active and stalled replication forks. Pol α is enriched at extending Okazaki fragments of active and stalled forks. In contrast, although Pol δ contacts the nascent lagging strands of active and stalled forks, it binds to only the matured (and not elongating) Okazaki fragments of stalled forks. Pol ε has a greater contact with the nascent ssDNA of leading strand on active forks compared with stalled forks. We propose that the configuration of DNA polymerases at stalled forks facilitate resumption of DNA synthesis after stress removal.

## Introduction

During eukaryotic genome duplication, replication stress is known to cause DNA synthesis to stall until the stress is alleviated. Replication stress includes lesions induced by endogenous and exogenous DNA-damaging agents, ribonucleotide mis-incorporation, and formation of secondary structures or DNA-RNA hybrids (1, 2). To better understand how genome integrity is maintained throughout replication stress, it is critical to determine how replisomes associate with DNA in active and stalled replication forks.

In budding yeast, DNA replication initiates at multiple sites, termed autonomously replicating sequences (ARSs) or replication origins. These origins are regulated temporally, with some origins firing early and others firing late in S phase of the cell cycle (3). In budding yeast, one of the primary responses to DNA replication stress is activation of the Mec1 and Rad53 kinase signaling cascade, a process equivalent to ataxia telangiectasia mutated– and Rad3-related (ATR) activation in human cells. Activated checkpoint kinases inhibit firing of late replication origins, maintain the stability of stalled replication forks, and help restart DNA synthesis.

DNA polymerases α, ε, and δ (Pol α, Pol ε, and Pol δ) are the main replicative DNA polymerases for eukaryotic nuclear genome, but other proteins are also involved in the process of DNA replication, including the Origin Recognition Complex (ORC) and replicative helicase minichromosome maintenance (MCM) proteins. In G1/S transition, MCM is activated through formation of the CMG complex (Cdc45, Mcm2-7, and GINS complex) and phosphorylation two kinases, CDK and DDK(4). Activated CMG helicase unwinds double-stranded DNA (dsDNA) at origins and recruits the single-stranded DNA (ssDNA)–binding protein RPA; RPA facilitates recruitment of Pol α, which synthesizes RNA primers followed by short DNA chain to initiate leading strands and Okazaki fragment synthesis. Pol ε and Pol δ extend Pol α’s products. In budding yeast, based on mutation bias introduced by Pol ε and δ mutants, it was proposed that Pol ε and δ are responsible for the synthesis of the leading and lagging strands, respectively (5–7). Recently, by mapping ribonucleotides introduced by DNA Polε and Polδ mutants genome wide in both budding yeast and fission yeast (8–11), it is deduced by that Polε and Polδ are involved in synthesis of leading and lagging strand, respectively. We have shown that Pol ε and Pol δ are enriched at nascent leading and lagging strands, respectively (12). However, a recent study of budding yeast suggest that Pol δ is involved in the synthesis of both leading and lagging strands and Pol ε is involved in DNA repair (13).

We and others have shown that proliferating cell nuclear antigen (PCNA), a processivity factor for Pol ε and Pol δ, is unloaded from lagging strands of stalled DNA replication forks and that this unloading is regulated by checkpoint kinases (12). Therefore, while replication proteins still associate with replisomes under stress so that DNA synthesis can resume once DNA replication stress is terminated, their contacts with DNA at stalled fork may differ from those with active forks. To analyze the interaction of DNA polymerases with replication forks in budding yeast, we evaluated the association of Pol α, Pol ε, and Pol δ with DNA in active and hydroxyurea (HU)-stalled replication forks by using chromatin immunoprecipitation plus strand-specific next-generation DNA sequencing (ChIP-ssSeq). Here, we report an in-depth analysis of protein ChIP-ssSeq datasets, which reveals distinct pattern of interaction of Pol δ and Pol ε with DNA at active and stalled replication forks. We suggest that these changes in contact with DNA directly or indirectly help maintain stability of stalled forks and facilitate the resumption of DNA synthesis after amelioration of replication stress.

## Methods

### Yeast strains

Yeast strains used in this study were derived from W303 (*leu2*-*3*, *112 ura3*-*1 his3*-*11*, *trp1*-*1*, *ade2*-*1 can1*-*100*). Genotypes are listed in Supplemental Table 1.

### ChIP-ssSeq Procedure

ChIP-ssSeq experiments were performed as described previously (12). Briefly, α factor was used to synchronize yeast cells at G1 (5 μg/mL and 50 ng/mL for wild-type *BAR* and *bar1* mutant strains, respectively). To analyze the association of proteins with active forks, G1-arrested cells were released into chilled (16°C) YPD medium containing 400 mg/L bromodeoxyuridine (BrdU), and samples were collected at different time points. We treated cells with HU, an inhibitor of ribonucleotide reductase, to deplete cells of dNTP. Thus, HU stalls the progression of DNA replication forks and inhibits the firing of late replication origins (14, 15). To analyze protein association with HU-stalled forks, cells were released into fresh medium containing 400 mg/L BrdU and 0.2M HU for 45 minutes. To perform ChIP, samples were incubated with 1% paraformaldehyde at 25°C for 20 minutes and then quenched with 0.125 M glycine for 5 minutes. Cells were lysed with glass beads, and the chromatin pellet was washed and sonicated to shear DNA to an average fragment size of about 200-400 bp. Sheared chromatin was immunoprecipitated with anti-FLAG antibody F1804 or anti-RPA antibody (gift of Dr Steven Brill). After extensive washing, cross-links of the immunoprecipitated chromatin were reversed. DNA was recovered with the Chelex-100 protocol (16). Recovered DNA was purified with a PCR purification kit (Qiagen). ChIP DNA was used to Q-PCR analysis (Supplemental Table 2) and were treated at 95 °C for 5 min before used for library preparation of ssDNA in accordance with previously published procedures (17).

### ChIP-ssSeq sequencing and analysis

The ssDNA libraries were sequenced using paired-end sequencing on Illumina Hi-Seq 2000 or 2500 machines. Reads were first mapped to the yeast genome (sacCer3) using Bowtie2 software (18). Consistent pair-end reads were chosen for subsequent analysis. We noted that after removal of duplicated reads, pair-end reads with the same ends were rarely detected in our samples even for Mcm6 ChIP-seq using G1 cells. This is likely due to the fact that chromatin was sheared by sonication and the ends were processed during library preparation. The genome-wide read coverage of Watson and Crick strands was calculated by BEDTools (19). The reads of the Watson and Crick strands were merged for peak calling by using MACS software (20).

We used our previously mapped DNA origins dataset for analysis (12). To calculate the average bias pattern, the average log_2_ ratios of sequencing reads of Watson strand over Crick strand surrounding 134 early replication origins (±10 Kb and ±30 Kb of HU-stalled and active forks, respectively) were calculated using a sliding window of 200-bp. The duplicate reads were excluded from calculation. These ratios were then normalized against the corresponding input to obtain the average bias pattern of ChIP-ssSeq. To analyze bias at individual origins, each peak region was separated into 4 quadrants: Watson strand at the left (WL) and right (WR) of an origin and Crick strand at the left (CL) and right (CR) of an origin. The number of sequence reads in each quadrant was counted. The binomial distribution was used to calculate the *P* value to determine whether sequence reads at leading strand (WL+CR) were different from those of the lagging strand (WR+CL) at each replication fork. The log_2_ ratio (log_2_ [(WL+CR)/(WR+CL)]) at each replication origin was calculated and used to determine whether a ChIP-ssSeq peak had a positive or negative strand bias.

Published dataset used in this study: Gene Expression Omnibus under accession number GSE52614.

## Results

### Rationale for analyzing replication proteins using ChIP-ssSeq

ChIP-sequencing (ChIP-seq) has been widely used to study the association pattern of a protein of interest with chromatin (21). Most ChIP-seq libraries are prepared using protocols that involve ligation of dsDNA, which often leads to loss of ssDNA and strand-specific information (Fig. 1A). During DNA replication, dsDNA is unwound to generate ssDNA, which serves as the template for DNA synthesis. In the process, replication proteins, including ssDNA-binding proteins RPA and DNA polymerases, may partially interact with the ssDNA or DNA-RNA hybrids. In addition, DNA replication forks consist of leading and lagging strand DNA synthesis, and strand-specific information helps elucidate how a protein interacts with forks.

**Figure 1.**
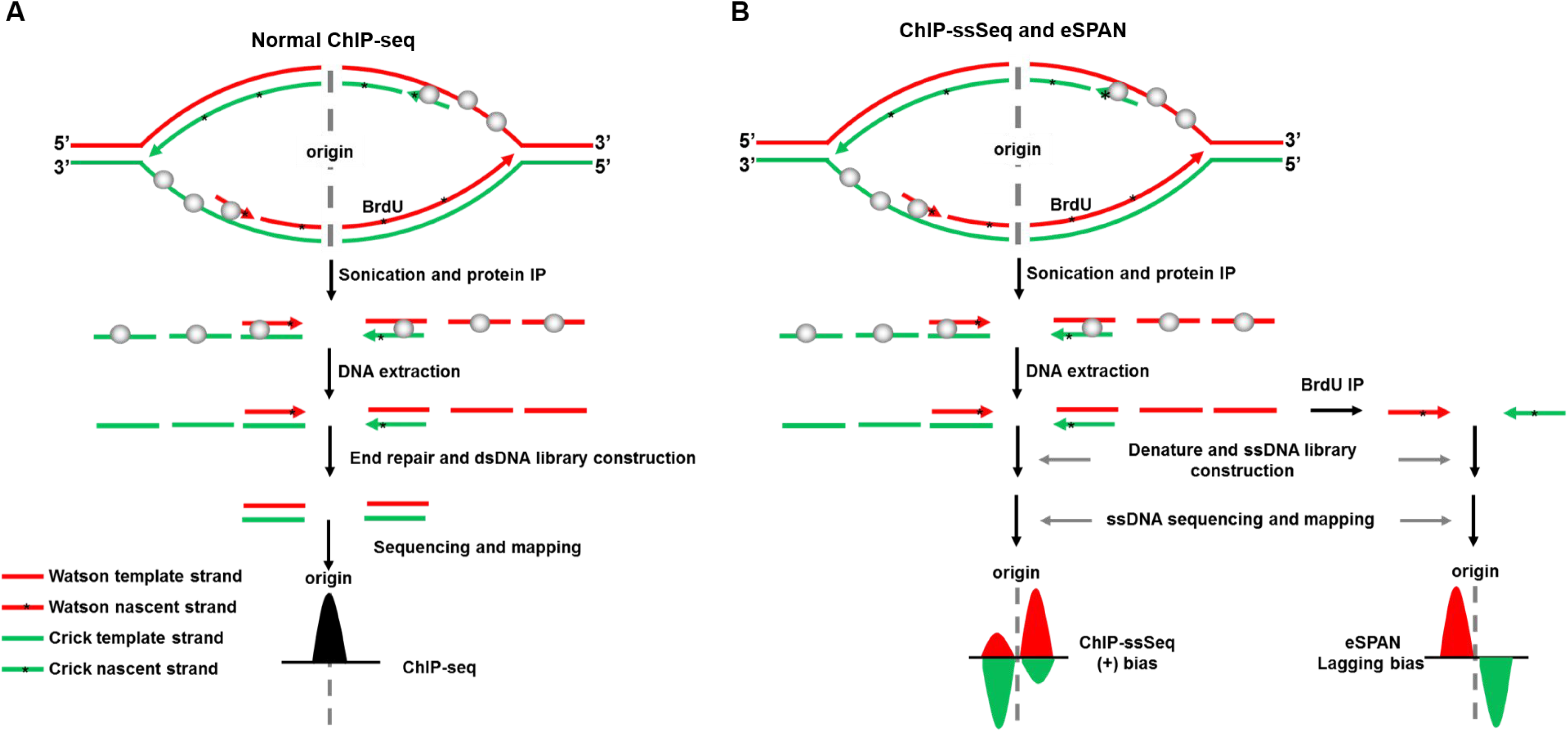
Schematic of the ChIP-ssSeq method used to detect the association of a protein with ssDNA. **(A)** The standard ChIP-seq process does not detect protein-ssDNA interactions. Here we use a protein that binds to both dsDNA and ssDNA as an example to elucidate the process. The upper shows a replication bubble. The first step is the protein ChIP. To prepare a library for sequencing, ChIPed DNA is extracted. After DNA end repair, only dsDNA is ligated with adaptors. Therefore, sequence reads contain location information for protein-dsDNA interactions. The target protein is shown as a gray ball. Black peak represent the DNA location. **(B)** The ChIP-ssSeq procedure preserves strand-specific information. The replication protein ChIP process is the same as the standard ChIP-seq. ChIP-ssSeq library preparation utilized a ssDNA ligase to ligate denatured ChIP ssDNA to a 3’ adaptor, which marks the same end of each ssDNA molecule. The second DNA strand is synthesized and extended with a 3’ complementary oligo. After end repair, the 5’ end is ligated to a 5’ end adaptor with T4 DNA ligase. After sequencing, the reads are mapped to the Watson and Crick strands of the yeast genome to determine the location of the target protein and strand-specific information. The red and green lines represent the Watson and Crick strands, respectively. ChIP-seq indicates chromatin immunoprecipitation and sequencing; ChIP-ssSeq, chromatin immunoprecipitation and strand-specific sequencing, dsDNA, double-stranded DNA; ssDNA, single-stranded DNA; ^∗^ represents nucleotide analog BrdU, which is incorporated into nascent DNA during DNA replication. As a comparison, the outline for eSPAN procedures is shown in B. The eSPAN procedure involves immunoprecipitation of protein-associated newly synthesized DNA marked with BrdU using antibodies against BrdU. Therefore, eSPAN detects the association of a protein with nascent DNA at DNA replication forks.

We previously reported development of the enrichment and sequencing protein–associated nascent DNA (eSPAN) method, which detects the association of a replication protein with nascent leading/lagging strand DNA (Fig.1B, right panel) (12). However, this method loses the information on how a protein interacts with ssDNA, which is prevalent at DNA replication forks. We also generated ChIP-ssSeq datasets during the process of obtaining eSPAN datasets. Briefly, protein ChIP DNA was denatured and ligated to the 3’ end of an adaptor (oligo) (Illumina) with an ssDNA ligase; ssDNA was then converted into dsDNA and ligated to a second adaptor (17, 22). The sequence reads were mapped to the Watson and Crick strands of the yeast genome (Fig. 1B). Since DNAs for a protein ChIP-ssSeq likely contain both template and nascent DNA (Fig.1B, left panel), ChIP-ssSeq will allow us to deduce how a DNA replication protein associates with single-stranded template DNA. As discussed and shown below, the ChIP-ssSeq and the eSPAN are two complementary methods, with each revealing unique information on the association of a protein at DNA replication forks.

### RPA ChIP-ssSeq shows that RPA is enriched at the lagging strand template

We first analyzed Rfa1 ChIP-ssSeq datasets to gain insight into how RPA associates with DNA replication forks. Briefly, yeast cells were arrested at G1 and then released into early S phase in the presence of HU for 45 minutes. Rfa1 (the large subunit of the RPA complex) ChIP was performed with G1 cells and early S-phase cells. Rfa1 was barely detectable at the replication origin (*ARS607*) or at a distal site (ARS607+8 kb, unreplicated region) at G1 (Supplemental Fig. 1A and 1B). In contrast, Rfa1 was enriched 10-fold at *ARS607* compared with the distal site (ARS607+8 kb) in the presence of HU (Supplemental Fig. 1B), indicating that RPA is recruited to DNA replication forks during S phase. Under these conditions, replication checkpoint kinase Rad53 is activated as shown by Western blot analysis of Rad53 (Supplemental Fig. 1C). In addition, the fact that late origins were not fired under these conditions also reflects the activation of Rad53 checkpoint kinase. Rfa1 ChIP-ssSeq peaks at *ARS510* and *ARS511* were asymmetric surrounding each origin (Fig. 2A), consistent with RPA binding to ssDNA and not to dsDNA. We note that a previous study shows that RPA binds asymmetrically to resected ssDNA in a double-strand break site (23).

**Figure 2.**
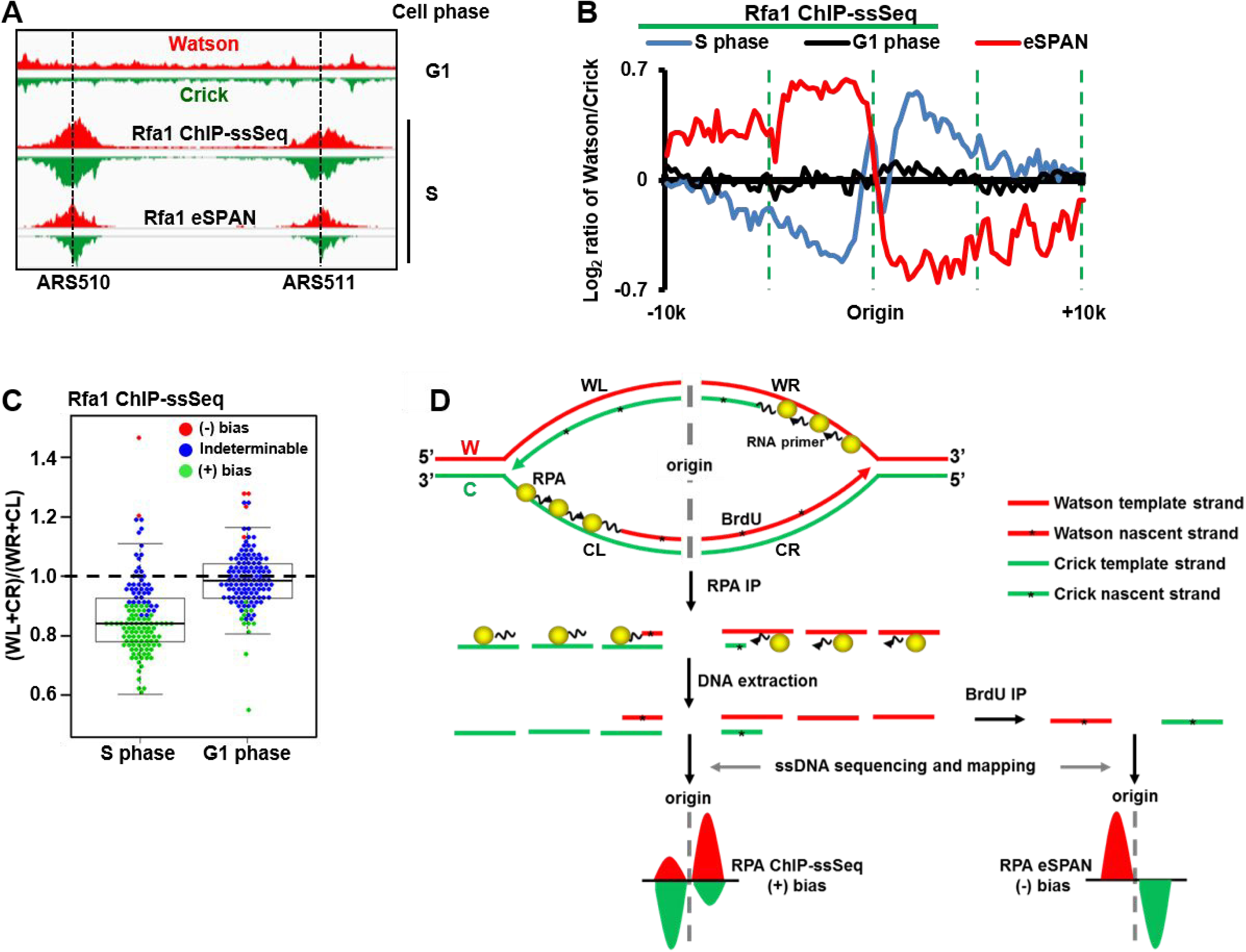
RPA preferentially binds to ssDNA on the lagging strand template. **(A)** A snapshot of RPA ChIP-ssSeq peaks at 2 early replication origins, *ARS510* and *ARS511.* Log-phase cells were synchronized to G1 with a factor and then released into 0.2M HU for 45 minutes. G1 and early S-phase cells were collected for RPA ChIP-ssSeq. The RPA ChIP was performed using antibodies against the FLAG epitope, which was fused to the C-terminus of Rfa1 (the large subunit of the RPA complex). Similar results were obtained by using an Rfa1-specific antibody (data not shown). The red and green regions represent normalized sequence read density of the Watson and Crick strands, respectively. Rfa1 eSPAN peaks are included for comparison (12). **(B)** The average pattern of Rfa1 ChIP-ssSeq peaks shows positive bias. The average log_2_ ratios of sequence reads of Watson and Crick strands at 134 early replication origins were calculated using a 200-bp sliding window and then normalized against input values to obtain the average bias pattern (blue). The lagging strand bias pattern of Rfa1 eSPAN peaks (12) (red) was used for comparison. **(C)** The dot-and-box plot shows the bias pattern of Rfa1 ChIP-ssSeq peaks at individual origins. The ratio of sequence reads of lagging to leading strands was calculated at each of 134 early replication origins; each dot represents one origin. The colors represent 3 bias patterns: red for positive bias (+); green for negative bias (−); and blue for indeterminable. **(D)** Schematic of a mechanism that accounts for the positive bias pattern of RPA based on ChIP-ssSeq analysis and the lagging strand bias pattern based on eSPAN analysis. The yellow ball represents the RPA complex. ^∗^ represents nucleotide analog BrdU, which is incorporated into nascent DNA during DNA replication. Please note that the eSPAN peak bias reflects whether a protein binds to nascent leading and lagging strand, whereas ChIP-ssSeq peak bias indicates that whether a protein binds to ssDNA or dsDNA.

To analyze Rfa1 ChIP-ssSeq results quantitatively at a genome-wide scale, we first calculated the average bias pattern, which is the average log_2_ ratio of sequencing reads of Watson strand over Crick strand using 200-bp sliding window surrounding 134 early replication origins. The average bias pattern of Rfa1 ChIP-ssSeq peaks indicated that on the right side of origin, RPA bound more to the Watson strand, whereas on the left side of origin, it bound more to the Crick strand (Fig. 2B). We categorized this finding as a positive (+) bias pattern to differentiate it from the leading-strand bias pattern revealed by the eSPAN method, which detects the association of a protein with newly synthesized DNA (12). As controls, the Rfa1 ChIP-ssSeq using G1 cells did not show any bias (Fig. 2B), suggesting that bias seen in early S phase reflects how RPA associates with DNA replication forks in the presence of HU. We also analyzed the bias pattern of Rfa1 ChIP-ssSeq peaks at each of the 134 individual replication origins (Fig. 2C). Rfa1 ChIP-ssSeq peaks showed (+) bias for most origins (n=89 [66%]), whereas the Rfa1 ChIP-ssSeq using G1 phase cells showed no bias for the majority of origins (n=119 [89%]). These results support the idea that RPA binds ssDNA of DNA replication forks stalled by HU.

While RPA is known to bind single-stranded template DNA, it may also contact nascent DNA at replication forks indirectly through protein-protein interactions. Indeed, RPA eSPAN reveals that RPA bind more to nascent lagging strands (12). Rfa1 ChIP-DNA contains the template strand and the nascent strand DNA. Two potential mechanisms account for the (+) bias pattern of Rfa1 ChIP-ssSeq peaks. First, (+) bias may indicate that more RPA binds to the lagging strand template than to the corresponding leading strand template (Fig. 2D). Second, RPA may bind more nascent leading strands than the corresponding nascent lagging strands. However, the later explanation contradicts the RPA eSPAN results outlined above (12) (Fig 2D). Based on our Rfa1 ChIP-ssSeq and Rfa1 eSPAN results, we suggest that more RPA binds lagging strand template than leading strand template of HU-stalled forks (Fig. 2D). The above RPA ChIP experiment is under HU condition. We also performed the RPA ChIP-ssSeq under normal condition. The results showed the same (+) bias pattern (Supplemental Fig. 1D-E), suggesting that more RPA are enriched at lagging strand template compared to leading strand template at both active and HU stalled forks. This explanation is consistent with the proposed model of RPA preferentially binding the lagging template strand to protect gaps between Okazaki fragments (24). To our knowledge, the result is the first experimental demonstration that more RPA binds lagging strand template than leading strand template. In addition to DNA replication, RPA is also involved in DNA repair process and activation of DNA replication checkpoint (25, 26).

### PCNA ChIP-ssSeq shows no strand bias at replication forks

We analyzed PCNA ChIP-ssSeq datasets obtained from cells cultured with or without HU. No obvious strand bias was observed from the analysis of the average bias pattern of all early replication origins or the analysis of the bias pattern of individual origins (Supplemental Fig. 2A-C). These results indicate that PCNA, a processivity factor of DNA polymerases that is loaded onto primer-template junctions, contacts dsDNA including both template and nascent DNA at active and HU-stalled replication forks.

### MCM ChIP-ssSeq shows no strand bias at stalled replication forks

Analysis of Mcm6 ChIP-ssSeq showed no significant strand bias of HU-stalled forks (Supplemental Fig. 2D-F), suggesting that the MCM helicase binds to dsDNA. Similar results were obtained for Mcm4 ChIP-ssSeq (Supplemental Fig. 2D-F). This observation seems to contradict the idea that the MCM complex travels along the leading strand (12, 27), and our eSPAN results showing that MCM associates preferentially with nascent leading strand DNA compared with nascent lagging strand DNA. One likely explanation for our ChIP-ssSeq results is that the MCM helicase, while encircling one leading template DNA strand, still makes indirect contact with another lagging template strand of HU-stalled forks. Indeed, it has been shown that MCM protein complex interacts with both Polε, which is enriched at leading strand, and Polα, which is enriched at lagging strands (28–31). For the rest of our studies, we focused on analysis on how three DNA polymerases associate with active and HU-stalled replication forks.

### Pol α ChIP-ssSeq indicates that Pol α preferentially binds to DNA-RNA hybrids at lagging strands of active and HU-stalled replication forks

Pol α was enriched at the early replication origin (*ARS607*) compared with the distal site (ARS607+8 kb) when cells were released from G1 to early S phase in the presence of HU (Supplemental Fig. 3A-B), consistent with the results that Polα associates with replicating DNA even in the presence of HU (32, 33). Inspection of Pol α ChIP-ssSeq at replication origins ARS510 and ARS511 showed that Pol α ChIP-ssSeq peaks showed a strong (+) bias pattern (Fig.3A and 3B). Analysis of the average bias pattern of 134 early replication origins confirmed that the (+) bias pattern of Pol α ChIP-ssSeq peaks at individual forks on a genome-wide scale (Fig. 3B), with 126 of 134 peaks (94%) showing (+) bias (Fig. 3C). We also determined how Pol α bound to active replication forks by performing ChIP-ssSeq using cells released into S phase without HU at a lower temperature (Fig. 3C and Supplemental Fig. 3C). The Pol α ChIP-ssSeq peaks also showed (+) bias based on the analysis of average bias pattern of early replication origins, as well as in the analysis of individual origins (Fig. 3C-E).

**Figure 3.**
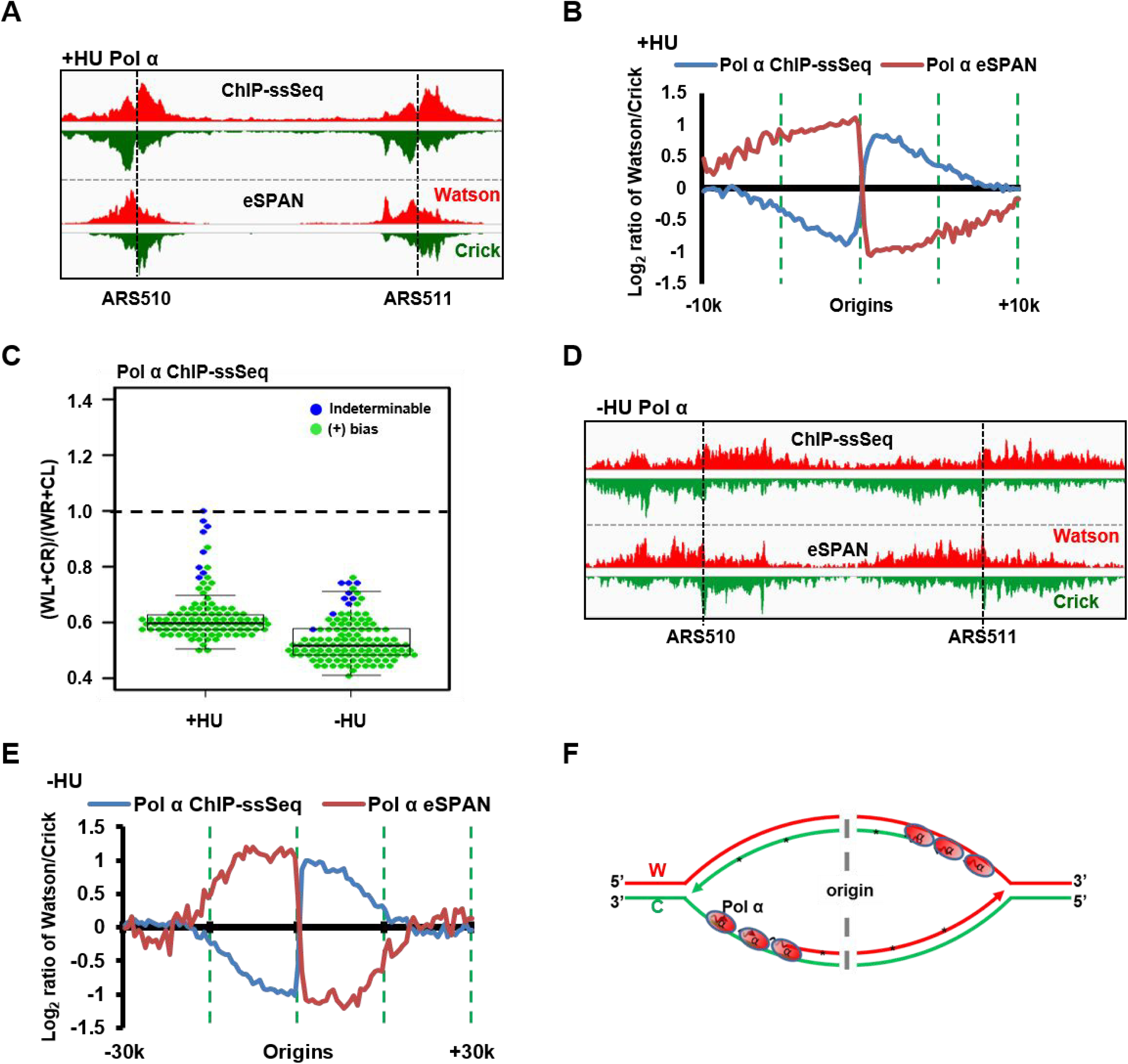
Pol α preferentially binds the single-stranded lagging template of both active and HU-stalled replication forks. **(A-C)** Pol α ChIP-ssSeq peaks at HU-stalled forks exhibit a positive (+) bias pattern. **(A)** Snapshot of Pol α ChIP-ssSeq and eSPAN peaks at *ARS510* and *ARS511.* The signals represent normalized sequence read density. The red and green represents the Watson and Crick strands, respectively. **(B)** Analysis of the average bias pattern of Pol α ChIP-ssSeq. Pol α eSPAN was used for comparison. **(C)** Analysis of bias pattern of Pol α ChIP-ssSeq of HU-stalled and active replication forks at individual origins (early replication origins only). **(D-E)** Pol α ChIP-ssSeq peaks at active forks show positive strand bias. G1-synchronized yeast cells were released into fresh medium at 16°C in the presence of BrdU for 72 minutes. **(D)** Snapshot of Pol α ChIP-ssSeq and eSPAN peaks at *ARS510* and *ARS511.* **(E)** Analysis of the average bias of Pol α ChIP-ssSeq peaks at active forks. f, Schematic of a mechanism that shows why Pol α ChIP-ssSeq has a positive strand bias pattern. Red line: Watson strand, Green line: Crick strand.

Pol α ChIPed DNA consists of the lagging strand template and newly synthesized RNA-DNA primer. The (+) bias pattern of Pol α ChIP-ssSeq peaks indicates that more Pol α binds to lagging strand template than to leading strand template of active and HU-stalled replication forks. Supporting this idea, the published Pol α eSPAN peaks indicate that Pol α physically binds more to nascent lagging strands than to leading strands at active and HU-stalled replication forks (12) (Fig. 3B, E-F). Because Pol α is involved in the synthesis of RNA and DNA primers, the (+) bias indicates that Pol α binds to initiating Okazaki fragments at HU-stalled forks and to the elongating Okazaki fragments of active replication forks (Fig. 3F).

### Pol δ is enriched at elongating Okazaki fragments of lagging strand template only of active replication forks

We next analyzed Pol δ (catalytic subunit) ChIP-ssSeq obtained using cells released from G1 arrest into early S phase in the presence of HU. Pol δ ChIP-PCR analysis showed that Pol δ was enriched at replication forks originating from *ARS607* comparing to the unreplicated distal site (ARS607+8 kb) (Supplemental Fig. 4A-B). Pol δ ChIP-ssSeq peaks at HU-stalled forks did not reveal any bias pattern based on analysis of the average bias of 134 forks from early replication origins or with analysis of individual origins (Fig. 4A-C), suggesting that Pol δ binds equally to Watson and Crick strands of HU-stalled replication forks. In contrast, Pol δ ChIP-ssSeq peaks at active replication forks without HU showed small, but consistent (+) bias in the analysis of the average bias pattern and of individual forks at all three-time points (Fig. 4B-C). The difference in Pol δ ChIP-ssSeq peak bias between active and HU-stalled forks was unlikely due to difference in input samples (Supplemental Fig. 4C). Thus, Pol δ differentially associates with DNA at active and HU-stalled replication forks

**Figure 4.**
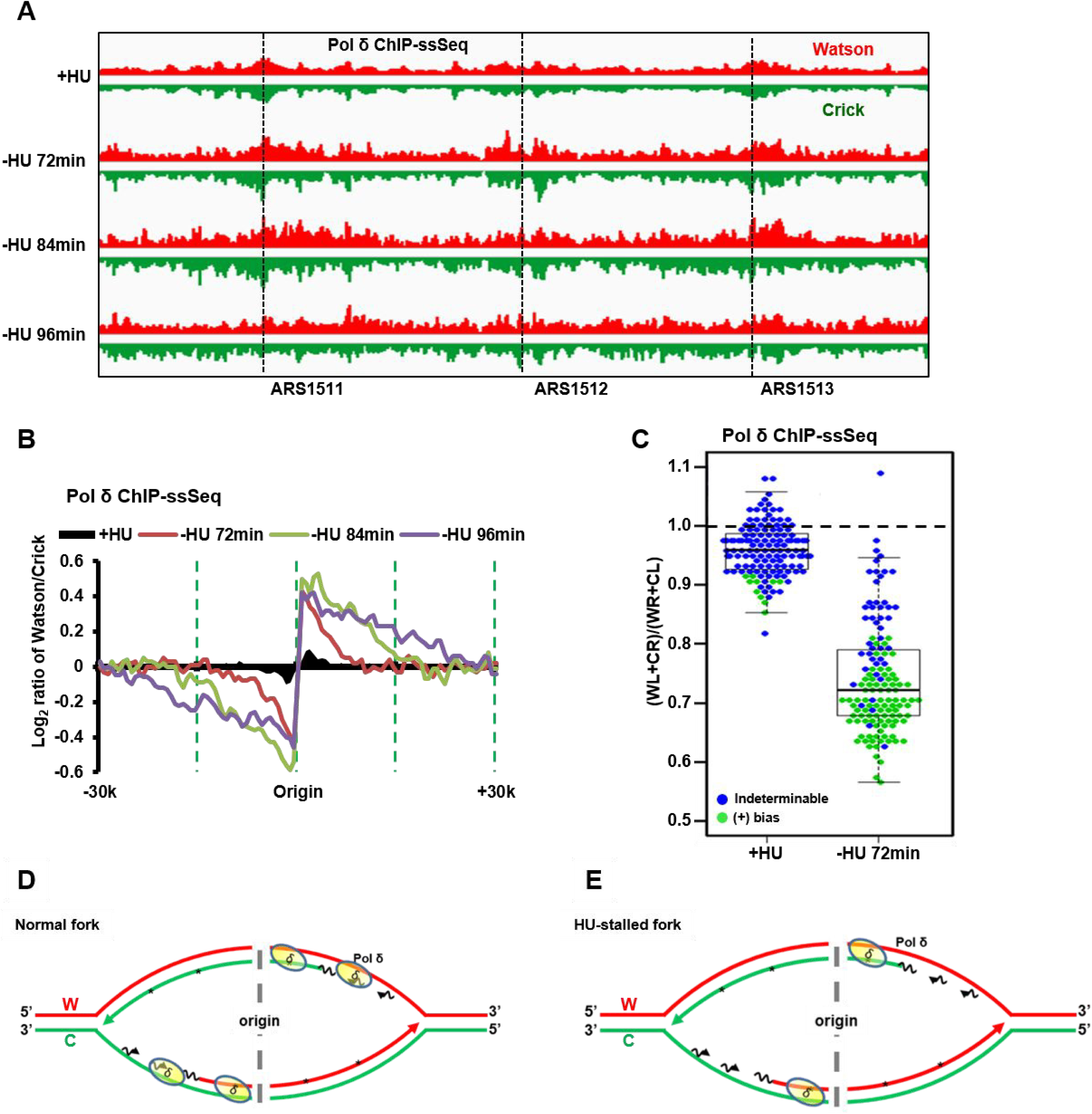
Pol δ binds dsDNA at HU-stalled replication forks and preferentially binds lagging-strand template ssDNA of active forks. **(A)** A snap shot of Polδ ChIP-ssSeq at three origins (*ARS1511*, *ARS1512* and *ARS1513*) with HU or three time points without HU. G1-synchronized yeast cells were released into fresh medium at 16°C without HU. Samples at the indicated time points (72 min, 84min and 96min) samples were collected for Pol δ ChIP-ssSeq of active forks. Sample was collected at 45 minutes after G1 release into HU at 30 °C for Pol δ ChIP-ssSeq at HU-stalled forks. Please note that Polδ ChIP efficiency was relatively low compared to Polε and Polα despite repeated attempts. **(B)** Pol δ ChIP-ssSeq peaks show positive bias at active forks and no bias at HU-stalled forks. The average bias pattern of ChIP-ssSeq peaks at 134 early replication origins is shown. The bias pattern indicated that Pol δ associates with dsDNA, but associates with ssDNA more frequently at active forks **(C)** Dot-and-box plot shows the bias pattern of Pol δ ChIP-ssSeq peaks at 134 individual early replication origins of HU-stalled and active forks. Each dot represents one origin. **(D-E)** Schematics show the Pol δ-DNA interaction at active forks **(D)** and HU-stalled forks **(E)**.

In principle, Pol δ binds both template and nascent DNA. Therefore, Pol δ ChIP-ssSeq peaks should show no bias at both active and HU-stalled forks. The eSPAN analysis of Pol δ indicates that Pol δ binds preferentially nascent lagging strand of HU-stalled and active replication forks (12) (Fig. 4B). We therefore deduce from the (+) bias pattern of Pol δ ChIP-ssSeq peaks that Pol δ associates with more lagging strand template of active forks, which most likely reflect that Pol δ can associates with newly initiated Okazaki fragments with only very short nascent RNA of active forks (Fig. 4D-E). In contrast, this mode of association of Pol δ is lost at HU-stalled forks, which provides an explanation for a lacking of bias of Pol δ ChIP-ssSeq peaks of HU-stalled forks. We have shown recently that the DNA polymerase clamp, PCNA, is unloaded from lagging strands of HU-stalled forks (12). PCNA is important for the activity of Pol δ, likely important for tethering Pol δ at DNA replication forks. Therefore, the unloading of PCNA from lagging strand of HU stalled forks may contribute to the loss of the association of Pol δ newly initiated Okazaki fragment at HU-stalled forks, whereas Pol α still binds.

### Pol ε-DNA interaction is different for active and the HU-stalled replication forks

After determining the association of Pol α and Pol δ with DNA, we next used ChIP-ssSeq to examine how Pol ε interacts with DNA. Pol ε ChIP-PCR analysis indicated that Pol ε bind to replicating DNA at HU-stalled replication forks (Supplemental Fig. 4A-B). Like Pol δ, Polε ChIP-ssSeq showed that Pol ε did not show significant bias at HU-stalled replication forks from early replication origins (Fig. 5A-C), indicating that Pol ε is cross-linked to dsDNA, including the template strand and nascent leading strand of HU-stalled replication forks. Remarkably, Pol ε ChIP-ssSeq showed (+) bias at actively replicating forks at all time points considered (72, 84, and 96 minutes after release from G1) (Fig. 5B). The bias pattern was detected at the majority of individual origins (Fig. 5C), suggesting that the Pol ε-DNA interaction at active forks differs from that at stalled forks.

**Figure 5.**
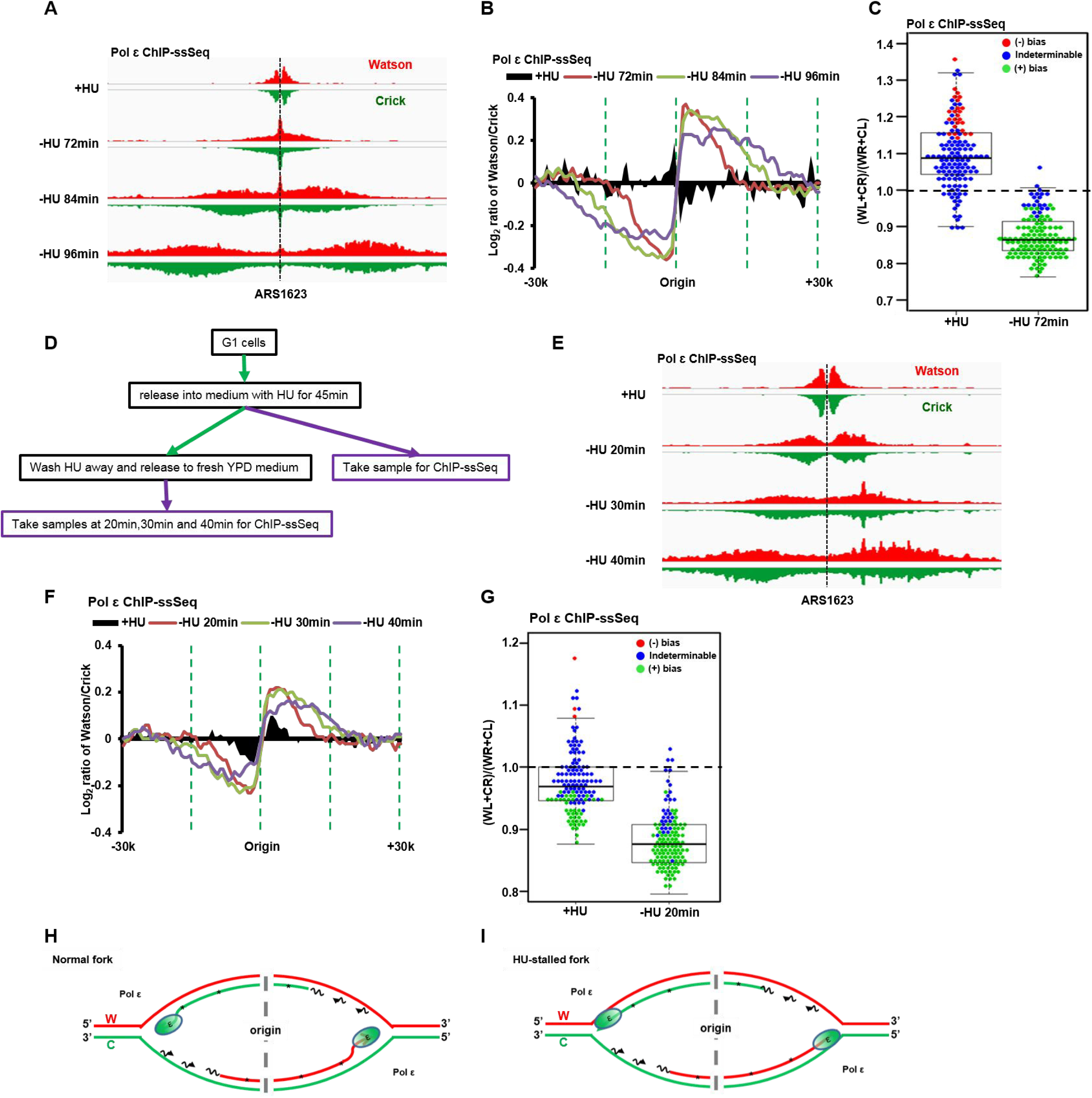
Pol ε binds dsDNA at HU-stalled replication forks and preferentially binds leading nascent ssDNA of active forks. **(A-C)** Pol ε ChIP-ssSeq peaks show a positive bias pattern at active forks and no bias at HU-stalled forks. The experiments were performed as described in Fig. 4. except that Pol ε ChIP-ssSeq was performed. **(A)** a snap shot of Pol ε ChIP-ssSeq peaks at *ARS1623.* **(B)** the average bias pattern of Pol ε ChIP-ssSeq peaks using a 200-bp sliding window(early replication origins only). **(C)** Dot-and-box plot shows the bias pattern of Pol ε ChIP-ssSeq peaks at 134 individual early origins. **(D-G)** The bias patterns of Pol ε ChIP-ssSeq for HU-stalled and active fork are different. The bias patterns indicate that Pol ε associate with dsDNA, but associates with ssDNA more frequently at active forks. **(D)** Flowchart of the experimental procedure. Yeast cells arrested in G1 were released into HU for 45 minutes. A fraction of cells were collected for Pol ε ChIP-ssSeq. The remaining cells were released into fresh media after removal of HU. Samples were collected at the indicated time points after release for Pol ε ChIP-ssSeq. **(E)** A snap shot of Pol ε ChIP-ssSeq peaks at *ARS1623* obtained using HU arrested or released cells. **(F)** Analysis of the bias pattern of Pol ε ChIP-ssSeq peaks at HU-stalled and active forks (3 time points after release). **(G)** Analysis of Pol ε ChIP-ssSeq peaks at individual origins at active and HU-stalled forks. **(H-I)**. Schematics showing Polε at active **(H)** and HU-stalled forks **(I)**. We propose that Polε binds to both template and nascent DNA of active forks, but with a higher frequency to nascent DNA than template DNA, which leads to the generation of Polε ChIP-ssSeq peak bias at active forks.

The above Pol ε ChIP-ssSeq analysis of HU-stalled and active replication forks were obtained from independent experiments and Pol ε ChIP-ssSeq bias is small. Therefore, we performed additional experiments to confirm that different association patterns of Pol ε with DNA changes in stalled vs active forks. Briefly, yeast cells were arrested in G1 with α factor and then released into HU-containing medium for 45 minutes. A fraction of cells were collected for Pol ε ChIP-ssSeq, and the remaining cells were released into fresh medium without HU. Samples were used to perform Pol ε ChIP-ssSeq at 3 time points after release from HU (20, 30, and 40 minutes) (Fig. 5D). Analysis of Pol ε ChIP-ssSeq datasets showed no bias pattern for peaks obtained using cells treated with HU, whereas peaks from cells after HU removal showed (+) bias (Fig 5E-G). We noticed that Polε ChIP-ssSeq at HU conditions shown in Fig. 5B and 5F appears to show opposite trend. This is likely due to the fact that Polε ChIP-ssSeq peaks at most origins showed indeterminable bias (no bias) and variations at a small number of origins contributes to the apparent changes in the insignificant bias pattern (compare Fig. 5C and Fig. 5G). Nonetheless, we observed very consistent results of Polε ChIP-ssSeq at active forks from each of the 3 time points of two independent experiments, supporting the idea that the association of Pol ε with DNA is altered when active forks become stall by HU treatment.

Once again, two potential models explain the (+) bias pattern of Pol ε ChIP-ssSeq peaks (Fig. 5H). Based on Polε eSPAN results (12) (Fig. 5B), Polε binds preferentially to leading strand. Therefore, it is possible that in addition to contact with leading strand DNA, Pol ε may also directly contact the lagging strand template during normal replication. This mechanism is unlikely because it is hard to put the Cdc45-MCM-GINS complex, which is known to associate with Pol ε on the leading strand (29), in front of Pol ε. Second, Pol ε may not contact the leading strand template tightly, binding only to nascent DNA on the leading strand of active forks (Fig. 5H). We suggest that this mode of interaction with leading nascent DNA facilitates its ability to proofread or repair mis-incorporated nucleotides by using its 3’-to-5’ exonuclease activity (34). At stalled fork, Pol ε may backtrack and associate with dsDNA including both template and nascent strands (Fig. 5I).

## Discussion

Our present study reveals several novel insights into the contacts of proteins with active and HU-stalled forks. First, we provide the experimental evidence that RPA are enriched at lagging strand template compared to the corresponding leading strand template, consistent with the replication model on the role of RPA in DNA replication. Second, we show that Polα associates with lagging strand template of both active and HU-stalled forks. Third, we show that both Polδ and Pol ε bind to HU-stalled forks differently from active forks. Specifically, Polδ binds to both initiating and elongating Okazaki fragments at active forks, and is likely lost/removed from initiating Okazaki fragments at HU-stalled forks where Polα remains to be present. Polδ likely backtracks and associtates dsDNA at HU-stalled forks. These results provide insight into how DNA synthesis can resume soon after removal of HU-induced replication stress.

### Advantages and limitations of ChIP-ssSeq method and its comparison with the eSPAN method

The library preparation of traditional ChIP-Seq includes steps for dsDNA repair and dsDNA ligation. During the sample preparation process, protein-bound ssDNA and strand-specific information is lost (Fig. 1A). Generally, this loss is not an issue because most proteins bind dsDNA. However, during DNA replication, dsDNA is transiently unwound into ssDNA. Therefore, determining whether a DNA replication protein binds to ssDNA will help elucidate its mode of action in DNA synthesis. Here, we used ChIP-ssSeq to gain insights into how DNA replication proteins bind to active and stalled replication forks. Since DNAs for protein ChIP-ssSeq potentially contain both template and nascent DNA, we analyzed ChIP-ssSeq peaks by calculating the average log2 ratio of sequence reads of Watson over Crick strand. If a protein contacts dsDNA including both template and nascent DNA, the ratio should be zero without any bias towards Watson or Crick strand. If a protein binds ssDNA, the average log2 ratio of sequence reads of Watson over Crick strand of ChIP-ssSeq peaks is not zero. Indeed, ssDNA-binding protein RPA ChIP-ssSeq peaks at both active and HU-stalled forks exhibit (+) bias, indicating that more RPA are present at lagging strand template than leading strand template, consistent with replication models of RPA in DNA replication.

We reported previously the development of the eSPAN method, which detects how a protein associates with newly synthesized DNA at DNA replication forks (12). Using this method, we detect the association of different DNA replication proteins with nascent leading or lagging strand of DNA replication forks. For instance, we observed that Polε and Polδ are enriched at nascent leading and lagging strand, respectively, consistent with their division of labor during DNA synthesis. Interestingly, we also show that RPA is enriched at lagging strand using eSPAN. This result appears to contradict the idea that RPA binds and stabilizes single-stranded template DNA. Using ChIP-ssSeq, we show that RPA is enriched at lagging strand template compared to leading strand template. One explanation for the apparent discrepancy for RPA ChIP-ssSeq and eSPAN results is that RPA binds preferentially to lagging strand template, but contacts nascent DNA indirectly, likely through other proteins. In this way, RPA eSPAN peaks show lagging strand bias. Similarly, the PCNA eSPAN results show that PCNA is enriched at nascent lagging strand DNA of active forks and nascent leading strand DNA at HU-stalled forks, suggesting that PCNA is unloaded from lagging strand of HU-stalled forks (12). However, PCNA ChIP-ssSeq peaks at HU-stalled forks show no bias pattern. One explanation is that PCNA contacts both template DNA and nascent DNA at HU-stalled forks, which gives rise to the no bias pattern of PCNA ChIP-ssSeq peaks. Therefore, the bias pattern of eSPAN peaks and ChIP-ssSeq peaks has a different meaning: the eSPAN peak bias indicates how a protein associates with nascent leading and lagging strands of DNA replication forks, whereas ChIP-ssSeq peak bias reflects how a protein binds to ssDNA and dsDNA. Together, these results comparing ChIP-ssSeq and eSPAN of different proteins including RPA, PCNA and MCM, PCNA and DNA polymerases (below) indicate that the ChIP-ssSeq and eSPAN methods provide complementary information on how a protein associates with DNA replication forks.

In theory, ChIP-ssSeq is suitable for studying DNA repair and RNA transcription when strand-specific information is needed. In fact, some reports indicated that similar ChIP-ssSeq approaches can study the DNA repair process (23, 35). These studies used either sticky end dsDNA adaptor ligation or intramolecular microhomology to generate libraries. We adopted the ssDNA library preparation method developed by Meyer et al (17), which was used to analyze highly damaged DNA from ancient human samples. The advantage of this method, compared with the 2 published ssDNA library preparation methods, is high efficiency. Meyer’s method can generate libraries from very low quantities of DNA (22), and it therefore is very suitable for constructing libraries from the low amount of DNA isolated by ChIP experiments. We expect that ChIP-ssSeq may also yield useful information for other processes besides DNA replication and repair. For instance, allele-specific DNA methylation is known to occur frequently in mammalian cells (36). A combination of library preparation methods with immunoprecipitation of methylated DNA will, in principle, be able to differentiate between methylated and unmethylated alleles. Future studies are needed to test this idea.

### Association of Pol α with active and HU-stalled replication forks

The Pol α-primase complex synthesizes primers for subsequent DNA synthesis by Pol δ and Pol ε, likely at the lagging and leading strands of DNA replication forks, respectively. Previously, we used the eSPAN method to show that Pol α-primase is enriched at the nascent lagging strand of the DNA replication fork; this finding was consistent with the classical replication models that require α Pol α-primase complex for each Okazaki fragment (37, 38). In this study, we used the Pol α ChIP-ssSeq method to show that Pol α also binds more to the lagging strand template during DNA replication, further supporting the role of Pol α–primase in the classical DNA replication model. Interestingly, we observed that Pol α–primase remains binding to the lagging strand template at HU-stalled replication forks. We suggest that the association of Pol α-primase complex with the template strand under this condition may facilitate resumption of the replication process soon after amelioration of DNA replication stress. In addition, this association may serve as the target of cell cycle checkpoint kinases that regulate arrest of DNA replication forks during stress. Consistent with this idea, previous work has shown that primase connects DNA replication to the DNA damage response (39).

### Altered association of Pol δ and Pol ε with DNA replication forks stalled by replication stress

Pol δ replicates both leading and lagging strands in the SV40 *in vitro* replication system (40–42). However, genetic evidence from the past decade supports Pol ε as the leading-strand replication enzyme in yeast (5, 6, 8, 43, 44) and Pol δ is responsible for replicating lagging-strand DNA. Recently, the division of labor between Pol ε and Pol δ in DNA synthesis has come into question with genetic analyses of mismatch repair–deficient DNA polymerase mutants (13). Therefore, several eukaryotic DNA replication models have been proposed (45). In every model, Pol ε is always physically linked with MCM helicase on the leading strand, regardless of whether it is the major active leading-strand DNA polymerase or just a repair enzyme. The result is fully compatible with our eSPAN data, indicating that Pol ε is enriched at the replicating leading strands. In contrast, Pol δ is enriched at the nascent lagging strands of DNA replication forks. We show here that Pol δ and Pol ε asymmetrically bind to DNA of active replication forks, suggesting that these two polymerases also bind to ssDNA, but not solely to dsDNA at active replication forks. In contrast, at HU-stalled forks, Pol δ and Pol ε predominantly were bound to dsDNA. We suggest that at HU-stalled forks, Pol ε may backtrack to contact dsDNA.

Previous study has shown that MCM localization can be displaced several hundred base pairs from the origin by transcription regulation (46). While it is possible that transcriptional alteration during HU block contributes to the lack of bias Pol ε and Pol δ ChIP-ssSeq peaks at HU-stalled forks, it is unlikely for the following reasons, First, we show that the Pol ε ChIP-ssSeq peak bias pattern reappears after we release cells from HU block to fresh media, suggesting that Pol ε bias is associated with active replication forks. Second, it is known that HU has no apparent effect on initiation of DNA replication at early replication origins based on studies from many laboratories. Moreover, the observation that transcription can shift MCM localization was made in *rat*1 mutant cells in which transcription termination was reduced, whereas at HU-stalled forks, we did not observe such dramatic alterations in MCM distribution (12).

We noticed the bias of Polδ and Polε ChIP-ssSeq peaks at active forks is small compared to that of Rfa1 or Polα. The small bias is not likely an artifact of calculation because we analyzed ChIP-ssSeq data sets using two different methods. First, we used 200 bp sliding window to calculate bias from either 10 or 30Kb surrounding each replication origins of HU-stalled and active forks, respectively. The trend of each data point of Polδ and Polε ChIP-ssSeq show that the bias, while small, is not random. Second, we also analyzed whether there exist bias of Polδ and Polε ChIP-ssSeq peaks at individual replication forks (Fig. 4C, Fig. 5C and Fig. 5G) and found that the bias, while small, is statistically significant. Unlike RPA that binds ssDNA and Polα that synthesizes primers for Polδ and Polε, most Polδ and Polε likely still contact dsDNA including template DNA and newly synthesized DNA. Therefore, it is not surprising that Polδ and Polε ChIP-ssSeq bias is smaller than RPA or Polα. In conclusion, the bias changes at HU-stalled forks and active forks of Polδ and Polε ChIP-ssSeq, while small, reflect the polymerase-DNA spatial contacts at active forks.

## Acknowledgement

We thank Dr. Oscar Aparicio for yeast strains and plasmids. We thank Dr. Steven Brill for the anti-RPA antibody and Dr. June Oshiro for editing our manuscript.

